# Escort-like somatic cells mediate early mouse fetal ovarian development but surface-derived Lgr5+ cells support primordial follicles

**DOI:** 10.1101/855346

**Authors:** Wanbao Niu, Allan C. Spradling

## Abstract

Ovarian murine somatic cells are essential to form first wave medullar follicles and second wave primordial follicles. Using single cell RNA sequencing we characterized the transcriptomes of both somatic and germline ovarian cells during fetal and early neonatal development. Wnt4-expressing somatic cells we term “escort-like cells (ELCs)” interact with incoming germ cells and early developing cysts of both sexes. In the medullar region, ELCs differentiate into the granulosa cells of fast-growing first wave follicles. In contrast, after E12.5, Lgr5+ pre-granulosa cells ingress from the ovarian surface epithelium and replace cortical escort-like cells. These surface-derived cells become the main population of granulosa cells supporting primordial follicles, and differ in transcription from ELC derivatives. Reflecting their different cellular origins, ablation of Lgr5+ cells at E16.5 using Lgr5-DTR-EGFP eliminates second wave follicles, but first wave follicles continue to develop normally and support fertility. Our findings provide striking evidence that somatic cell behavior supporting germline cyst development in mice and Drosophila has been evolutionarily conserved.

## Introduction

The basic outlines of somatic cell development in the mammalian fetal ovary are well understood in rodents (Hirshfield, 1991; McLaren, 1991; Edson et al. 2009; Rotgers et a. 2018). In mouse, the coelomic epithelium (CE) forms on the ventral side of the mesonephros beginning at about E9.5, thickens, proliferates, and begins to express characteristic genes. Primordial germ cells (PGCs) reach the gonad about E10.5 where they proliferate mitotically to form cysts that partially fragment and aggregate together into cell “nests” (Pepling and Spradling, 1998; Mork et al. 2012b; Lei and Spradling, 2013). Female development requires Wnt4/Rspo1/β-catenin signaling (Vainio et al. 1999; Chassot et al. 2008; Maatouk et al. 2013) during this period, but the specific cells utilizing these pathways are not fully delineated. CE proliferation contributes to at least two distinct populations of pregranulosa cells as well as interstitial cells and steroid hormone producing cells, some of which are proposed to arise via Notch dependent asymmetric divisions (Lin et al. 2017). Eventually, two “waves” of follicles are produced; a first wave that develops rapidly in the medullar region of the ovary and a second wave derived from the primordial follicle pool in the cortex that provides fertility throughout life (Byskov et al,. 1997; Zheng et al. 2014). However, much remains to be learned about the lineage origins and transcriptomes of the somatic granulosa cells making up these two subpopulations of follicles.

Two populations of ovarian somatic support cells are known to interact with germ cells in cysts and in developing follicles. The first population arises by at least E11.5 from an unknown source, and may comprise bipotential precursors of either Sertoli or granulosa cells (McLaren, 1991; Mork et al. 2012a). Starting at E12.5 and continuing past E14.5, Foxl2-expression turns on in a subset of existing somatic cells, and lineage labeling showed that these cells give rise to some of the granulosa cells exclusively within first wave follicles (Mork et al. 2012a; Zheng et al. 2014). The second pregranulosa cell population is derived from Lgr5+ progenitors in the ovarian surface epithelium that migrate into the ovarian cortex (Ng et al. 2014; Rastetter et a. 2014) and generate granulosa cells on 2nd wave follicles (Mork et al. 2012a; Zheng et al. 2014). A third somatic cell type, ovarian steroidogenic thecal cells, differentiate shortly after birth from Gli1-expressing ovarian mesenchymal cells in a process dependent on hedgehog signaling from granulosa cells (Liu et al. 2015). Despite this progress, the cellular origins, interactions and gene expression programs of the granulosa cells surrounding first and second wave follicles remain imperfectly understood.

Critical insights into germ cell and follicle development have also come from genetic and physiological studies (reviewed in Handel and Schimenti, 2010; Rotgers, 2018). Genes responding to the meiotic inducer retinoic acid (RA) (Bowles et al. 2006), and to its key target Stra8, have been characterized (Soh et al. 2015; Kojima et al. 2019). Recently, mammalian and mouse germ cell development has been further analyzed using single cell RNA sequence (scRNAseq) analysis, especially in the male germ line (reviewed by Suzuki et al. 2019). Fetal mouse gonadal somatic cells of both sexes were purified and analyzed by scRNAseq to better understand early sex differentiation (Stevant et al. 2019). However, much remains to be learned about the genetic programs of female meiotic germ cells, the initial bipotential somatic cells, and later ovarian somatic cells that support both primary and secondary waves of follicle formation. A powerful adjunct to scRNAseq for the analysis of such questions is the ability to reconstruct developmental trajectories (Butler et al., 2018) and validation by lineage tracing.

Evolutionary conservation provides another source of insight into ovarian follicle development. In both mouse and Drosophila, primordial germ cells migrate to the gonadal primordium (Bendel-Stenzel et al. 1998; Richardson and Lehmann, 2010), and oocytes differentiate within interconnected cysts of meiotic germ cells with the assistance of nurse cells (Telfer, 1975; Lei and Spradling, 2016). In mice cysts are built using five rounds of synchronous division as well as limited breakage (Pepling and Spradling, 1998; Lei and Spradling, 2013). Inactivation of Wnt4/Rspo1 in mouse ovarian somatic cells results in germ cell losses and a partial male sex transformation (Vainio et al. 1999; Chassot et al. 2008; Maatouk et al. 2013). In Drosophila, early germ cells associate with somatic escort cells that express Wnts (Morris and Spradling, 2011; Kirilly et al. 2011; Wang et al., 2015a). Disruption of Wnt signaling in escort cells upregulates BMP signaling and interferes with germ cell development and survival (Mottier-Pavie et al. 2016; Wang and Page-McCaw, 2018). Escort cells are displaced from germ cells in mature ovaries by migrating follicle cells which then form a granulosa cell-like epithelial monolayer that mediates subsequent follicle development (Margolis and Spradling, 1995; Nystul and Spradling, 2007). Thus, there is evidence for the involvement of two types of supporting somatic cells during both mouse and Drosophila folliculogenesis, but whether there is any correspondence between the somatic ovarian cells of these two species remains unknown.

Here we profile cells generally in the neonatal mouse ovary using scRNAseq, including multiple somatic cell types that provide further insight into ovarian follicular development. Our analysis of germ cells confirms and extends information from previous studies (Soh et al. 2015; Wang et al., 2015b; Kojima et al. 2019). We describe a common initial population of “escort like cells” preferentially expressing Wnt4, Wnt6 and Bmp2, that include squamous cells contacting all forming germ cell cysts as previously observed by electron microscopy (Pepling and Spradling, 2001). Lineage tracing shows that ELCs on medullar follicles differentiate into granulosa cells, and support first wave follicle development. However, the ELCs in the cortical region are replaced by ingressing Lgr5-positive pregranulosa cells, such that primordial follicles develop with the exclusive support of surface-derived granulosa cells. This behavior strongly resembles the replacement of non-dividing Drosophila escort cells around cysts by migrating follicle cells in mature ovaries. Thus, our genetic and cellular findings support the view that the somatic as well as the germline program of follicle formation has been conserved in evolution.

## Results

### Generation of a single-cell transcriptional atlas of perinatal ovarian development

To investigate cellular diversification during early folliculogenesis in mice, we performed single cell RNA-seq on developing perinatal ovaries at E12.5 (sex determination), E14.5 (meiotic entry), E18.5 and P1 (cyst breakdown and oocyte selection), and P5 (primordial follicles) (see Table S1). The selected time points span key events during perinatal ovarian development. After a meticulous dissection and trypsin incubation, ovaries were dissociated into single-cell suspensions. 24,228 dissociated cells were subsequently captured, loaded onto oil droplets, and used for cDNA library construction, deep sequencing, and cluster analysis (Figure 1A; see Methods). Expression information on an average of 2,700 different genes was recovered from each cell.

**Figure 1.**
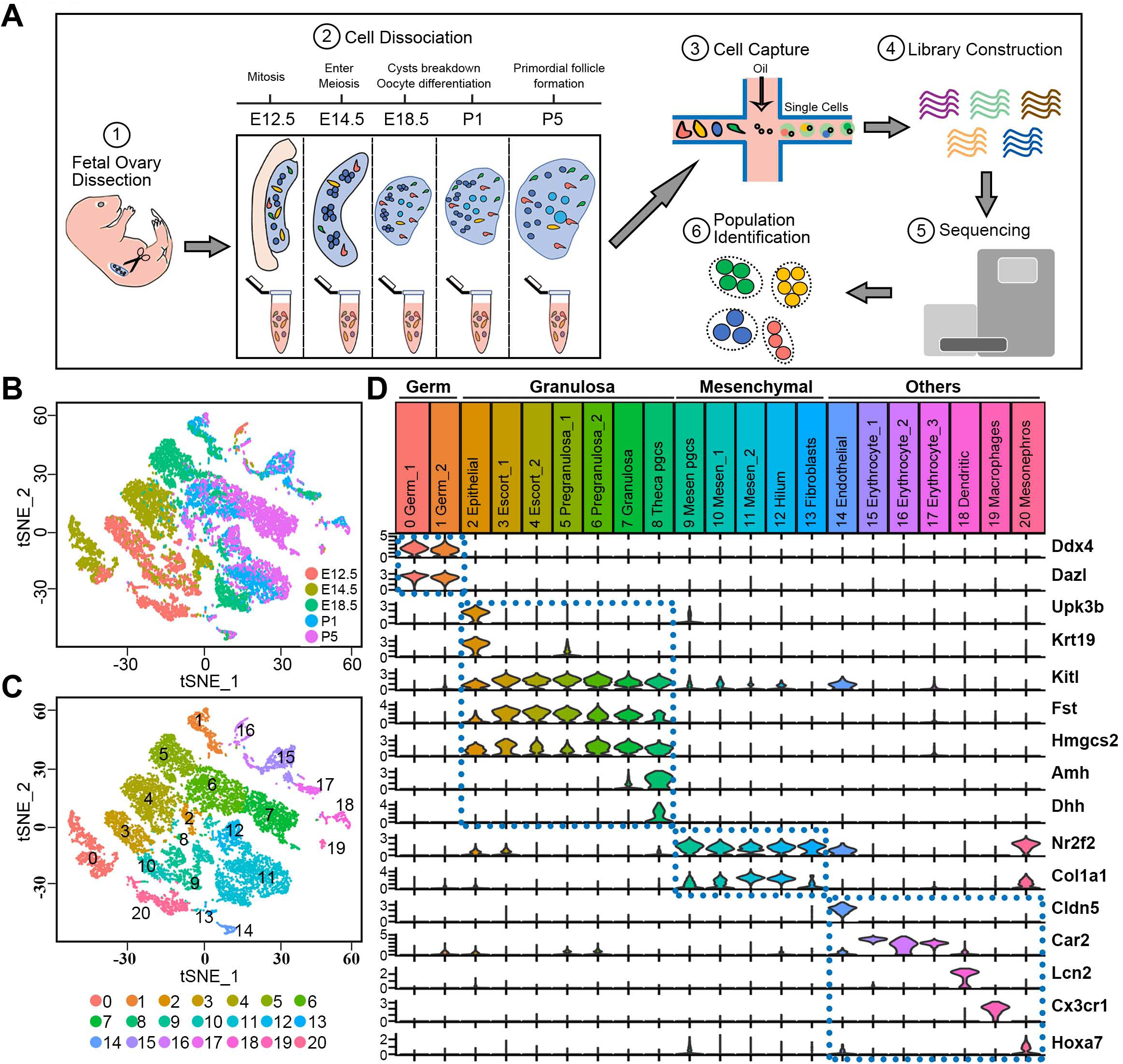
Single-cell transcriptome landscape of fetal ovarian development. (A) Schematic experimental workflow using the 10x Genomics Chromium platform followed by clustering using Seraut (Butler et al. 2018). (B) Two-dimensional visualization of single-cell clusters using tSNE colored by developmental time from E12.5 to P5. (C) Two-dimensional visualization of single-cell clusters using tSNE colored by 21 identified cell types/groups (numbers). (D) Summary of marker gene expression in cell clusters: Clusters 0-20 were sub-divided into four subclasses based on gene expression (dashed boxes). Below, violin plots below show marker gene expression in each cluster. y-axis scale: a normalized UMI/cell scale for each gene to facilitate display.

The datasets generated from the different time points were analyzed jointly using the following strategy. Transcript counts were first normalized, log2 transformed, aligned, and integrated as described (Butler et al., 2018). Using a t-distributed stochastic neighbor embedding (tSNE) analysis, we arranged the integrated datasets in temporal order (Figure 1B), and clearly identified 21 clusters (Figure 1C), classified as four major categories (Figure 1D). These were germ cells (cluster 0, 1) with Ddx4 and Dazl expression (Lin et al., 2008), granulosa cells (clusters 2-8) with Kitl, Fst and Hmgcs2 expression (Jones and Pepling, 2013; Wang et al., 2015c); mesenchymal cells (clusters 9-13) with Nr2f2 and Col1a1 expression (Ku et al., 2006; Rastetter et al., 2014); and others (clusters 14-20). This last category included endothelial cells (cluster 14) that express Cldn5 (Choi et al., 2012), three erythrocyte clusters (15, 16, 17) that uniquely express Car2 (Lewis et al., 1988), dendritic cells (cluster 18) with high Lcn2 expression (Mucha et al., 2011), macrophages (cluster 19) uniquely expressing Cx3cr1 (Medina-Contreras et al., 2011), and mesonephritic cells (cluster 20) that highly expressed Hoxa7 (Ng et al., 2016).

By immunofluorescence staining of ovary sections from different developmental stages using cluster predominant genes, we validated germline, mesenchymal and endothelial cell types that had been identified based on their transcriptomes (Figure 1D), and further analyzed granulosa-related cells in the perinatal ovaries. Co-staining for Nr2f2, Foxl2, and Ddx4 showed that Nr2f2-expressing cells and Foxl2-expressing cells are mutually exclusive both in E12.5 and E18.5 ovaries (Figure S1A). At E12.5, very few of the cells adjacent to germ cell cysts expressed Foxl2 whereas expression was widespread in these cells at E18.5, consistent with previous observations. At E18.5, the Nr2f2-positive mesenchymal cells align in rows that have been likened to cords. Staining mesenchymal cells located between groups of cysts with Col1a1 further highlighted these ovarian subdomains, each of which contains several cysts or primordial follicles in E14.5 and P2 ovaries (Figure S1B).

### Fine scale analysis of the germ cell meiotic transcriptome

We choose germ cells, whose gene expression during meiosis has been well studied, to investigate computational methods to further refine and extend our analysis. We selected only the germ population and re-performed tSNE analysis (Figure 2A). Early stage germ cells (E12.5, E14.5), which are completing mitotic divisions and just entering meiosis, were mapped on the left side of the tSNE plot, whereas later stage germ cells (E18.5, P1, and P5), which had finished synapsis and arrested at diplotene/dictyate, localized to right side of the plot largely in temporal order (Figure 2B).

**Figure 2.**
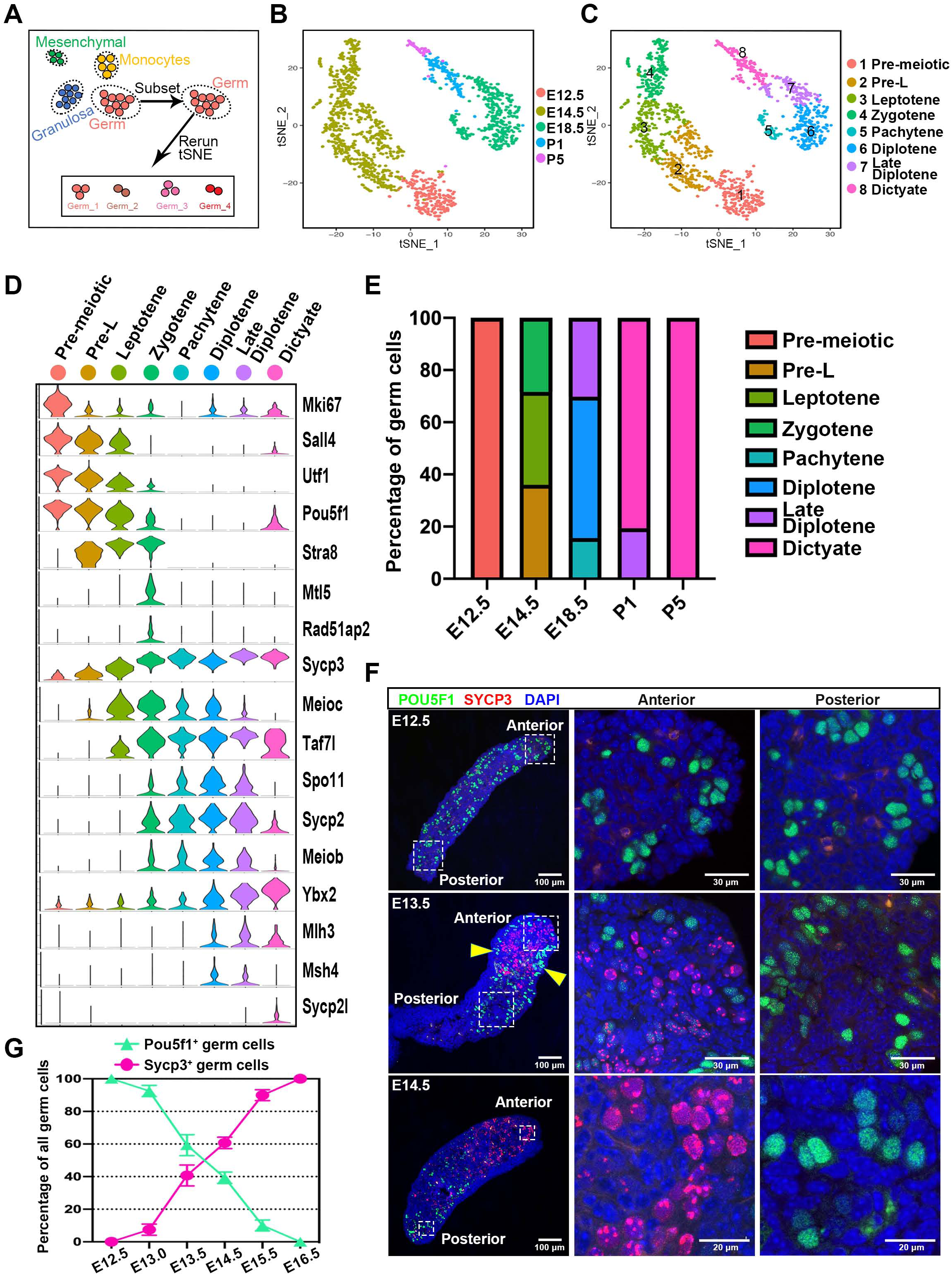
Dynamic gene expression patterns of mouse female germ cells. (A) Germ cells extracted at each time point combined and re-analyzed. (B) Two-dimensional visualization of clusters from all germ cells using tSNE. Cells colored stage of sequencing from E12.5 to P5. (C) Two-dimensional visualization of integrated germ cell clusters using tSNE. Cells colored by eight inferred developmental stage (see key for stage names). (D) Multi-violin plot of selected meiosis-related gene expression during the eight developmental stages. y-axis scale: same as Fig. 1D. (E) Cell distribution among the eight stages at each time point. (F) Immunofluorescence of E12.5, E13.5 and E14.5 ovaries co-stained for Pou5f1 and Sycp3, showing anterior-posterior expression gradient. (G) Quantification of germ cells expressing Pou5f1^+^ or Sycp3^+^ during fetal ovary development.

The germ population could then be computationally subdivided into eight transcriptionally distinct successive germ clusters, whose gene expression was found to match the known temporal pattern of gene expression during meiosis (Figure 2C, D; Table S2). Pre-meiotic germ cells (Fig. 2D, cluster 1) express the mitotic marker Mki67 and pluripotency markers (such as Sall4, Utf1 and Pou5f1). Pre-leptotene cells (cluster 2) still express Mki67 at very low levels, retain high levels of Sall4, Utf1 and Pou5f1, but turn on the key retinoid acid (RA) signaling pathway target Stra8 (Anderson et al., 2008) and upregulate the cohesin subunit Rec8, a marker of meiotic DNA replication (Dokshin et al. 2013). Leptotene cells (cluster 3) express pluripotency genes at reduced levels and Stra8 at higher levels than cluster2. They also express Sycp3 and Taf7l, and turn on Meioc, which plays a critical role in the leptotene/zygotene transition (Abby et al., 2016). Zygotene cells (cluster 4) now retain very little Sall4, Utf1 or Pou5f1 expression, but uniquely harbor Mtl5 and Rad51ap2, as well as a collection of meiotic genes including Sycp2 and Spo11, which are required to initiate meiotic recombination (Kovalenko et al., 2006; Yang et al., 2006). Clusters 6 and 7 differentially express the diplotene markers Ybx2, Mlh3, and Msh4 (Lipkin et al., 2002; Santucci-Darmanin et al., 2000; Wang et al., 2015b). Cluster 6 expresses higher Meioc but lower Ybx2 and Mlh3 than cluster 7, indicating that cluster 6 cells have just entered diplotene, while cluster 7 cells are near the diplotene/dictyate transition. Finally, dictyate cells (cluster 8) specifically expressed Sycp2l, which is only found in dictyate stage oocytes and regulates the survival of primordial oocytes (Zhou et al., 2015). Our analyses identified the expression profiles of more than 3,000 genes that varied substantially across the 8 identified meiotic substages (Figure S2; Table S2).

We next examined the relative proportion of each meiotic cluster contributing to the dataset at each developmental time point (Figure 2E). As expected, all germ cells scored as prior to meiosis at E12.5. Their developmental heterogeneity at E14.5 (pre-leptotene, 36.1%), leptotene (35.5%) and zygotene (28.4%) reflects the 24hr variation in time over which PGCs arrive at the gonad and begin developing. Starting at E18.5, most germ cells are at diplotene stages (pachytene 15.7%, diplotene, 54.1%; late diplotene, 30.2%), and by P1, most have arrested at dictyate (80.6%).

Immunofluorescence staining using Pou5f1 and Sycp3 antibodies in the early fetal ovaries validated the known spatial asynchrony of meiotic progression shown in the above cluster analysis, (Figure 2F). As expected, at E12.5, when the female germ cells undergo rapid division and form germline cysts, all germ cells exhibited strong staining with Pou5f1 but no staining with Sycp3. By E13.5, germ cells in the anterior region have begun to express Sycp3 while the germ cells at the posterior continue to show high Pou5f1 staining. It should be noted that germ cells located at the surface in the anterior surface still express Pou5f1 but not Sycp3 (arrowheads), indicating that there is also a temporal difference in meiotic timing between the cortex and deeper layers. At E14.5, the percentage of Sycp3+ germ cells increased to 60.7%. These inverse trends continued until Pou5f1 expression completely disappeared at E16.5 (Figure 2F, 2G). Thus, our analysis of germline gene expression identified the same meiotic stages across the multiple time points, despite temporal variation in when progenitors arrive at the gonad and spatial variation in germ cell development along the anterior-posterior ovarian axis.

### Escort-like cells initially associate with mouse germline cysts

We next investigated the somatic cells most likely to interact with germ cells. We removed the germ cells, mesenchymal cells and others, and reanalyzed the cells in the granulosa subclass using tSNE (Figure 3A). This identified fourteen granulosa subclusters, which as before were plotted by temporal order (Figure 3B) and by individual cluster type (Figure 3C). Known markers were used to identify subgroups of epithelial (Upk3b and Krt19), pre-granulosa (Lgr5 and Gng13), and granulosa (Hmgcs2) cells, leaving one major subgroup that we termed “escort-like cells (ELCs)” (Figure 3C, escort1-3; Table S3) after the Drosophila somatic cells that first interact with germ cells.

**Figure 3.**
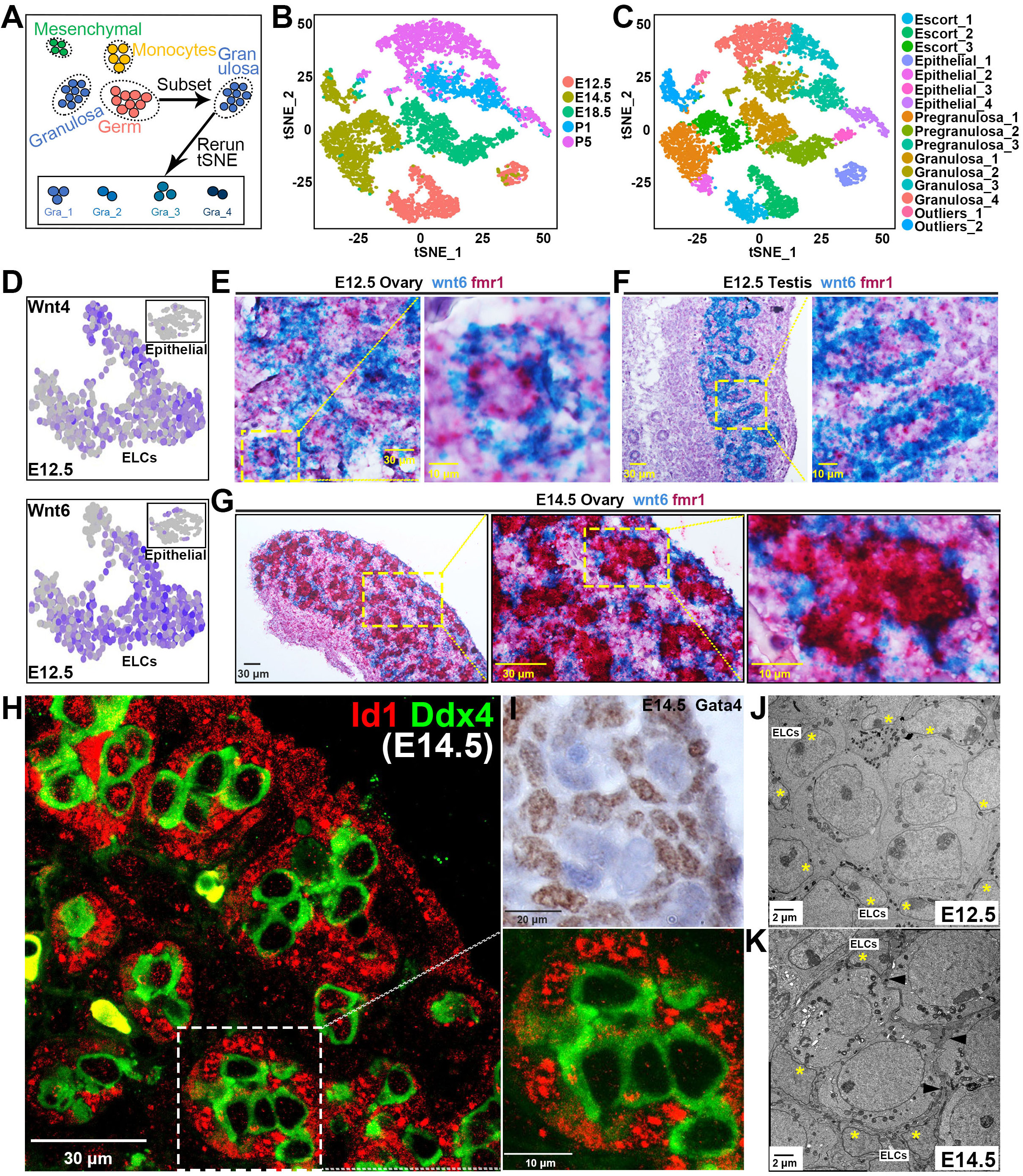
Identification and cellular localization of escort-like cells. (A) Granulosa cell sub-populations extracted, combined and re-analyzed. (B) Two-dimensional visualization of clusters from all “granulosa” subgroup cells using tSNE. Cell colored by embryonic time points from E12.5 to P5. (C) Two-dimensional visualization of granulosa cell clusters using tSNE. (D) Expression of Wnt4 and Wnt6 of escort-like cells, color indicates level of expression. (E-G) *In situ* hybridization (ISH) analysis shows Wnt6 (blue) and Fmr1 (red) mRNA expression in the E12.5 ovary (E), E12.5 testis (F), and E14.5 ovary (G). (H) Cellular localization of Id1 in E14.5 ovaries. Ovaries were stained for Id1, and the oocyte marker DDX4 at E14.5 by immunofluorescence. (I) Cellular localization of Gata4 in E14.5 ovaries by immunohistochemistry. J) Electron micrograph of E12.5 ovary showing part of a germline cyst (center) surrounded by ELCs (yellow asterisks). K) Electron micrograph of E14.5 ovary showing part of a germline cyst surrounded by ELCs (yellow asterisks). Squamous membranes of ELCs surrounding the germ cells are indicated by arrowheads. Scale bars are indicated.

ELCs are the only granulosa-like cells in E12.5 ovaries, and their transcriptome further suggests that they play critical roles in early germ cell development (Table S3). ELCs express Wnt ligands, including Wn4 and Wnt6, at higher levels than other E12.5 cells (Figure 3D; Figure S3A), suggesting that ELCs mediate Wnt-dependent early ovarian functions such as germ cell maintenance and sex determination (Vainio et al. 1999; Chassot et al. 2008). In addition, ELCs expressed Kitl, Rspo1,Wt1, Lhx9, Emx2 and granulosa cell markers at higher levels than other E12.5 cells, showing their likely identity as bipotential somatic precursors. Like these precursors (Mork et al. 2012a), ELCs appear to be non-mitotic, as they express only low levels of the mitotic marker Mki67, and high levels of Cdkn1b1 encoding a p27 cell cycle kinase inhibitor.

We used double in situ hybridization to investigate the cellular localization of ELCs in early gonads. Cells expressing Wnt6 mRNA were detected adjacent to germ cell nests (marked by Fmr1 expression) in the E12.5 ovary (Figure 3E). A similar pattern was observed in E12.5 testis where many nest/cysts are encircled by Wnt6-positive cells (Figure 3F). These observations argue that ELCs associate closely with germ cells in early ovaries, and include the somatic cells seen to surround developing germ cell cysts in electron micrographs of E12.5 ovaries (Figure 3J, asterisks). At E14.5, Wnt6-expressing cells continued to adjoin germline nest/cyst structures (Figure 3G). In EMs at E14.5, squamous somatic cells are seen to wrap each cyst (Fig 3K, arrowheads, asterisks).

The ELC transcriptome also provided evidence that these cells are active in Bmp signaling at E12.5. BMP2, but not other BMPs, as well as the Bmp target genes Id1, Id2, Id3, and Gata4 are expressed (Figure S3B). Immunofluorescence studies showed that cells positive for Id1 tightly wrap germline cysts/nests in the E14.5 ovary (Figure 3H). Staining E14.5 ovaries with anti-GATA4 antibodies gave an identical pattern (Figure 3I). These experiments indicate that ELCs express high levels of BMP target genes, and reinforce evidence that they tightly contact individual cysts/nests in early ovary.

### Cortical escort-like cells are replaced by Lgr5-positive pre-granulosa cells

A second major population of pregranulosa cells is generated from Lgr5-expressing progenitors in the ovarian surface epithelium (Mork et al. 2012a; Ng et al. 2014; Rastetter et a. 2014; Zheng et al. 2014; Lin et al. 2017). These cells begin to invade the ovarian cortex at E12.5 and continue to be produced past E14.5. Levels of both Wnt4 and Id1, which are highly expressed in ELCs, dramatically decrease after E18.5 (Figure 4A, B). In contrast, the pre-granulosa cell marker genes Gng13 and Lgr5 trended upward from E12.5 to E18.5 before falling at P1 (Figure 4C, D). These changes suggest that ELCs gradually turn over in concert with the arrival of surface derived pregranulosa cells beginning after E12.5.

**Figure 4.**
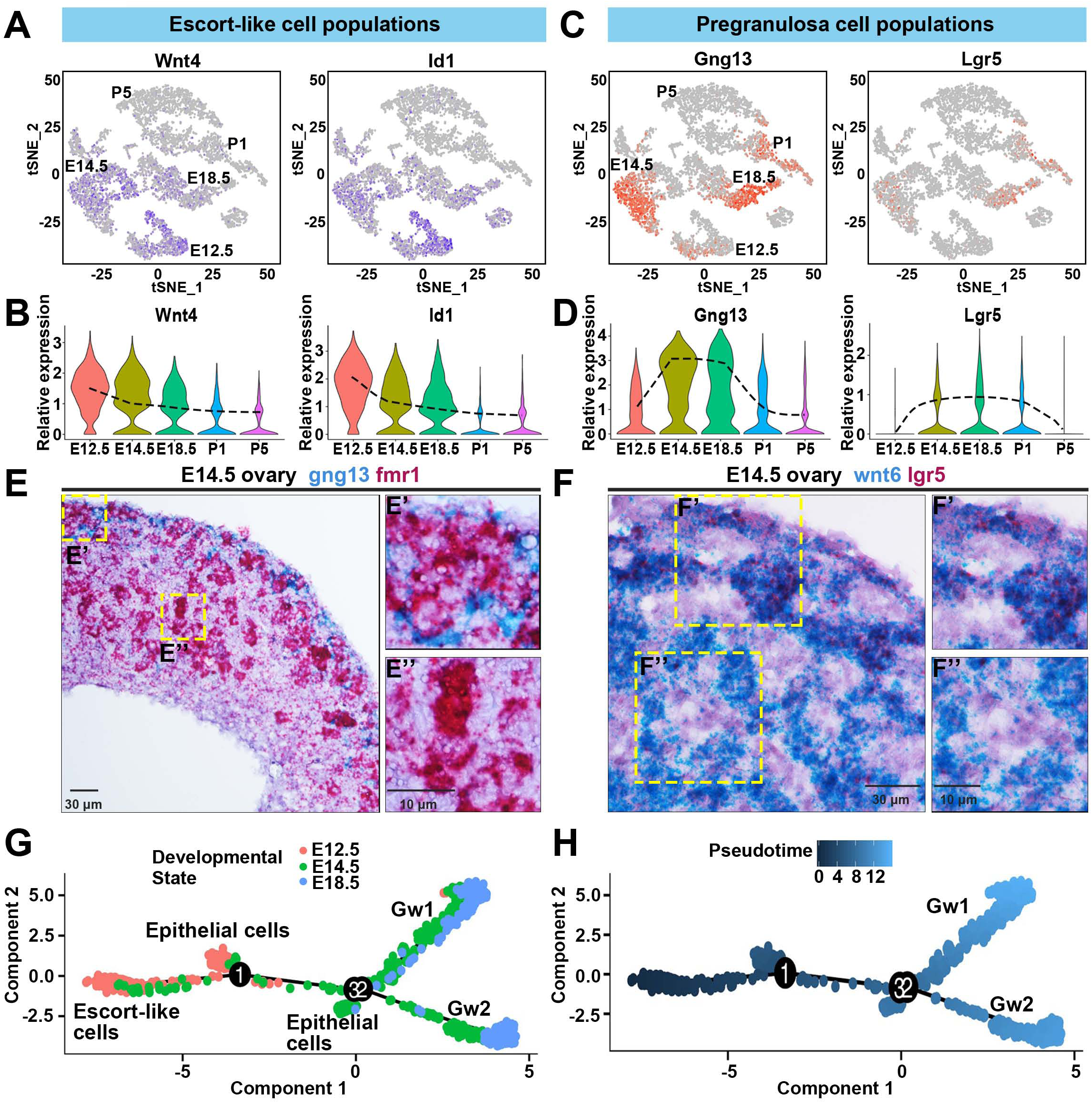
Escort-like cells are replaced by pregranulosa cells. (A) Expression of Wnt4 and Id1 of escort-like cells, color indicates level of expression. (B) The violin plot showing relative expression of Wnt4 and Id1 of granulosa population from ovaries at E12.5, E14.5, E18.5, P1, and P5. The dashed line indicates the expression trend of Wnt4 and Id1. (C) Expression of Gng13 and Lgr5 of surface-derived pregranulosa cells, color indicates level of expression. (D) The violin plot showing relative expression of Gng13 and Lgr5 of granulosa population from ovaries at E12.5, E14.5, E18.5, P1, and P5. The dashed line indicates the expression trend of Gng13 and Lgr5. (E) *In situ* hybridization (ISH) analysis shows Gng13 (blue) and Fmr1 (red) mRNA expression in the E14.5 ovary. Gng13-expressing pregranulosa cells are confined to the cortical (E’) but absent from the medullar (E”) subregions (yellow boxes). (F) In situ hybridization analysis shows Wnt6 (blue) and Lgr5 (red) mRNA expression in the E14.5 ovary. Lgr5^+^ pregranulosa cells were detected in the cortical (F’) but not in deeper (F”) subregions (yellow boxes). (G-H) Differentiation trajectory of E12.5, E14.5, and E18.5 ovary beginning with escort-like cells and epithelial cells and splitting into two branches (Gw1, Gw2) constructed by Monocle. Cell colored by the developmental state (G) or pseudotime (H).

We took several approaches to learning about the timing and fate of surface derived Lgr5+ pregranulosa cells after E12.5. We carried out in situ hybridization to localize cells expressing pre-granulosa cell marker genes such as Gng13 and Lgr5, and for ELC markers like Wnt6. At E14.5, Gng13 and Lgr5 both were expressed strongly in the ovarian surface epithelium (Figure 4E, F), and were detected in cells that had entered cortical regions near the surface and were now located adjacent to germ cell clusters (Figure 4E’, 4F’). Germline cysts located deeper in the ovary and in the medullar region were still surrounded by cells that did not express detectable levels of Gng13 or Lgr5 (Figure 4E”, 4F”) but were Wnt6-positive suggesting they were residual ELCs or cells derived from them (Figure 4F”). These data suggest that the cortical Gng13^+^ Lgr5^+^ pregranulosa cells represent surface epithelium-derived cells that have migrated inward and replaced ELCs in cysts at the ovarian cortex, but few if any of these cells have reached the inner cortex or medulla region by E14.5.

To gain further insight into the developmental fates of ELCs and surface derived pregranulosa cells, we carried out developmental trajectory analysis of these populations (Butler et al. 2018). We analyzed all epithelial, escort-like, pregranulosa and granulosa cells from E12.5, E14.5 and E18.5. This yielded a curve that connected E12.5 escort-like cells and epithelial cells with two distinct populations of E18.5 pregranulosa cells (Figure 4G, 4H). Based on comparison of gene expression differences (Figure S4A,B), we believe that the two populations correspond to pregranulosa cells on wave1 follicles (Gw1) or on wave two follicles (Gw2), and that they derive from escort-like cells or surface epithelial cells, respectively. For example, genes such as Map1b, Tuba1a and Greb1 are expressed at significantly higher levels in ELCs and in Gw1 cells (Figure S4A). In contrast, known gonad epithelial genes including Lgr5, Gng13 and Lhx9 are selectively expressed in Gw2 pregranulosa cells (Figure S4B). Many lipoprotein metabolism genes (such as Apoc1, Gpc3) and peptidase inhibitor marker genes (such as Cst8, Cst12) are also preferentially expressed in epithelial cells and Gw2 pregranulosa cells. We continued the pseudotime analyses of Gw1 and Gw2 cell classes using granulosa cells from P1 and P5 to provide a broader picture of the transcriptome development of these two subclasses of pregranulosa/granulosa cells (Figure S4; Table S4). This analysis strongly confirmed our identification of the two groups, because at P5, when some first wave follicles have already developed further than wave 2 follicles, and reached the primary follicle stage, Gw1 cells preferentially expressed multiple genes associated with later follicle development including Amh, (ratio Gw1/Gw2: 62.5), Esr2 (38.8), and Nr5a2 (14.8) (Durlinger et al. 1999; Chakravarthi et al. 2018, Meinsohn et al. 2017).

### ELC lineage tracing validates ELC replacement in the cortex but not the medulla

We validated the trajectory analysis using lineage marking of progenitors of the two major somatic cell populations. Since ELCs actively express axin2 (Figure S3A), we lineage labeled early ELCs using Axin2-CreERT2 mice. Axin2^CreERT2/+^ mice were crossed to Rosa26-YFP reporter mice, and pregnant females received tamoxifen at E10.5. Ovaries were analyzed at E12.5, E15.5, E19.5 and P21 (Figure 5A). As expected, at E12.5, ELCs were extensively labeled in both the cortex and medulla of the ovary, and were seen to contact virtually all germ cell cysts (Fig. 5B). By E15.5, however, the number of labeled cells in the cortical (but not the medullar) region was reduced. By E19.5 and again at P21, labeled cells were found exclusively in the medullar region, indicating that cortical ELCs have been fully replaced and turned over, rather than undergoing transdifferentiation (Fig. 5B). Figure 5C summarizes the loss of labeled ELCs from the cortex and their retention in the medulla. These experiments show that ELCs are initially found in association with all germ cell cysts, but are lost as surface derived cells move into the cortical region. ELCs remain in the medullar region and contribute to wave 1 follicles.

**Figure 5.**
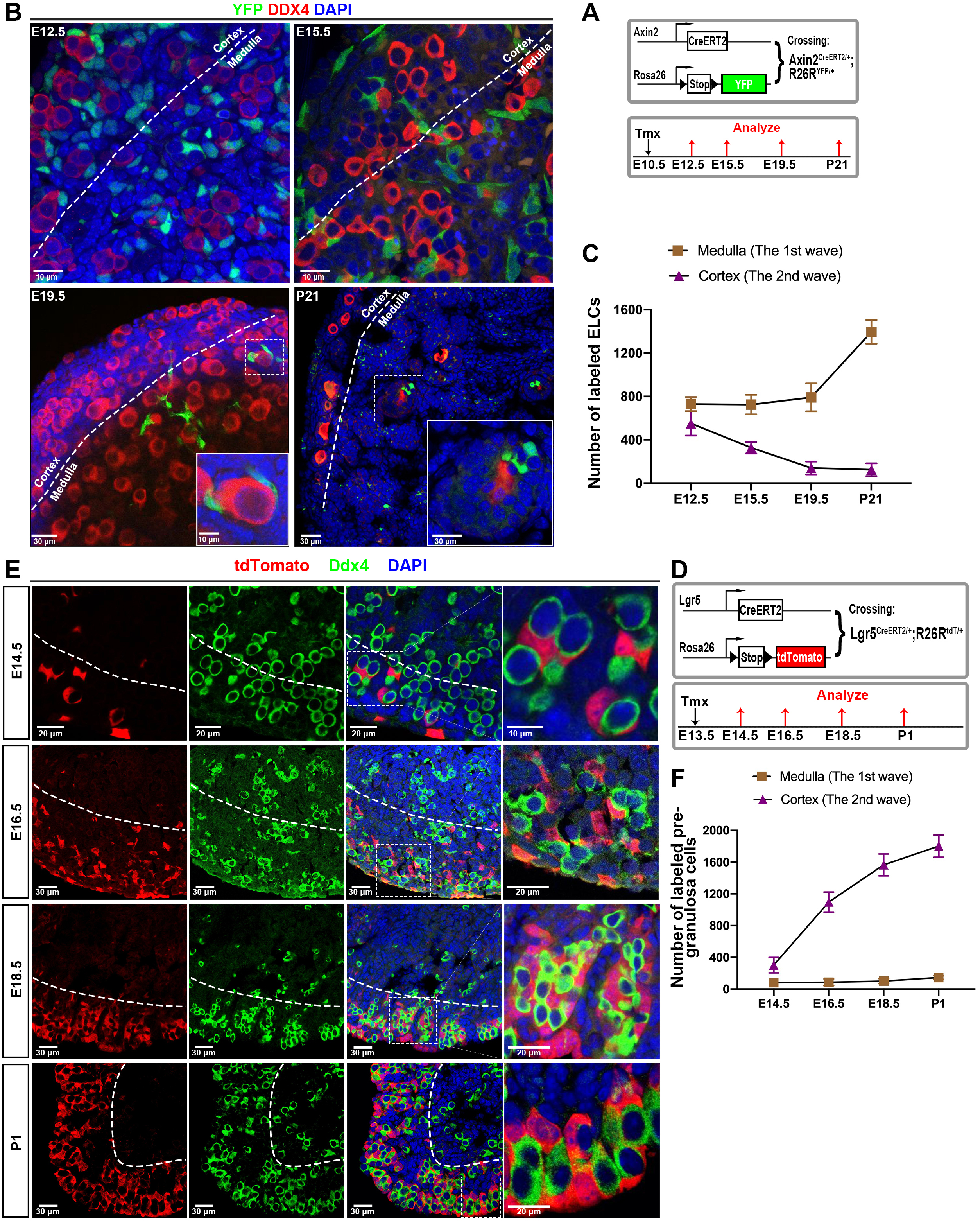
Lineage tracing validates and reveals the kinetics of cortical ELC replacement by Lgr5+ pregranulosa cells. (A) Schematic of ELC lineage tracing strategy. Mice containing Axin2^CreERT2/+^ mice and R26R^YFP/YFP^ received Tamoxifen (TM) at E10.5, and ovaries were collected at E12.5, E15.5, E19.5 and P21 for histological analysis. (B) Lineage tracing of Axin2+ escort-like cell progeny for E19.5 and P21 ovary demonstrates that Axin2+ escort-like cells mainly contribute to the first wave follicles. (C) YFP-labeled ELCs were quantitated in the cortex and medulla regions from ovarian sections at each time point, as shown in (B). (D) Schematic of the surface granulosa cell tracing strategy. Mice containing Lgr5^CreERT2/+^ and R26R^tdT/tdT^ reporter received Tamoxifen (TM) at E13.5 to activate red fluorescent protein (tdTomato) expression, and ovaries were collected at E14.5, E16.5, E18.5 and P1 for histological analysis. (E) Relatively few tdTomato positive pregranulosa cells had entered the cortical region by E14.5 but those present associated with germ cells. The number of cortical tdTomato+ cells increased significantly at E16.5, E18.5 and P1, while very few cells entered the medullar region (F). Dashed lines in B and E show the boundary of the cortical and medullar regions.

### Kinetics of Lgr5+ pregranulosa cell replacement of cortical ELCs

We further investigated the timing of Lgr5+ pregranulosa cell association with cortical follicles using linage marking. Lgr5^CreERT2/+^ mice were crossed to Rosa26-tdTomato reporter mice, and we administered TM to pregnant females at E13.5. Ovaries from offspring at 14.5, 16.5, 18.5 and P1 were then collected and analyzed (Figure 5D). Since dividing Lgr5+ cells are essentially confined to the ovarian surface at E13.5, this protocol is expected to label newly generated Lgr5^+^ pregranulosa cell progenitors and reveal their subsequent behavior.

When we analyzed mice at E14.5 whose ovarian cells had been labeled in this manner at E13.5, several results were clear (Figure 5E, F). The great majority of the tdTomato^+^ cells (clonally related to the Lgr5^+^ pregranulosa cells labeled at E13.5) were observed in the cortical region contacting germ cells. At E16.5, the number of tdTomato^+^ cells in the cortex increased greatly, while the small number of medullar cells showed little change. This trend continued at E18.5 and P1, by which time cortical germ cells appeared to be almost entirely surrounded by tdTomato^+^ cells (Figure 5E, F). The few remaining unlabeled cells were probably Lgr5^+^ cells that had not become marked, since ELC lineage marking showed these cells were no longer present. Consistent with the view that Lgr5 cells do not migrate from the cortex into the medulla, tdT^+^ cells remained at very low levels the medullary region. The few exceptions involved cells close to the cortical-medullary border.

We used this same marking system to examine Lgr5+ daughters labeled at P1 (Figure S5A). This allowed us to address whether continuing cell division at the ovarian surface produces new pregranulosa cells after birth. Our experiments showed that although many tdT-labeled cells were detected at both P2 and P5 at the ovarian surface, none of these cells had migrated into the ovarian cortex (Figure S5B,C). Thus, surface Lgr5+ cells continue to divide at P1, but cortical pregranulosa cells are no longer generated. A model summarizing somatic cell behavior during first and second wave follicle formation is shown in Figure 6A.

**Figure 6.**
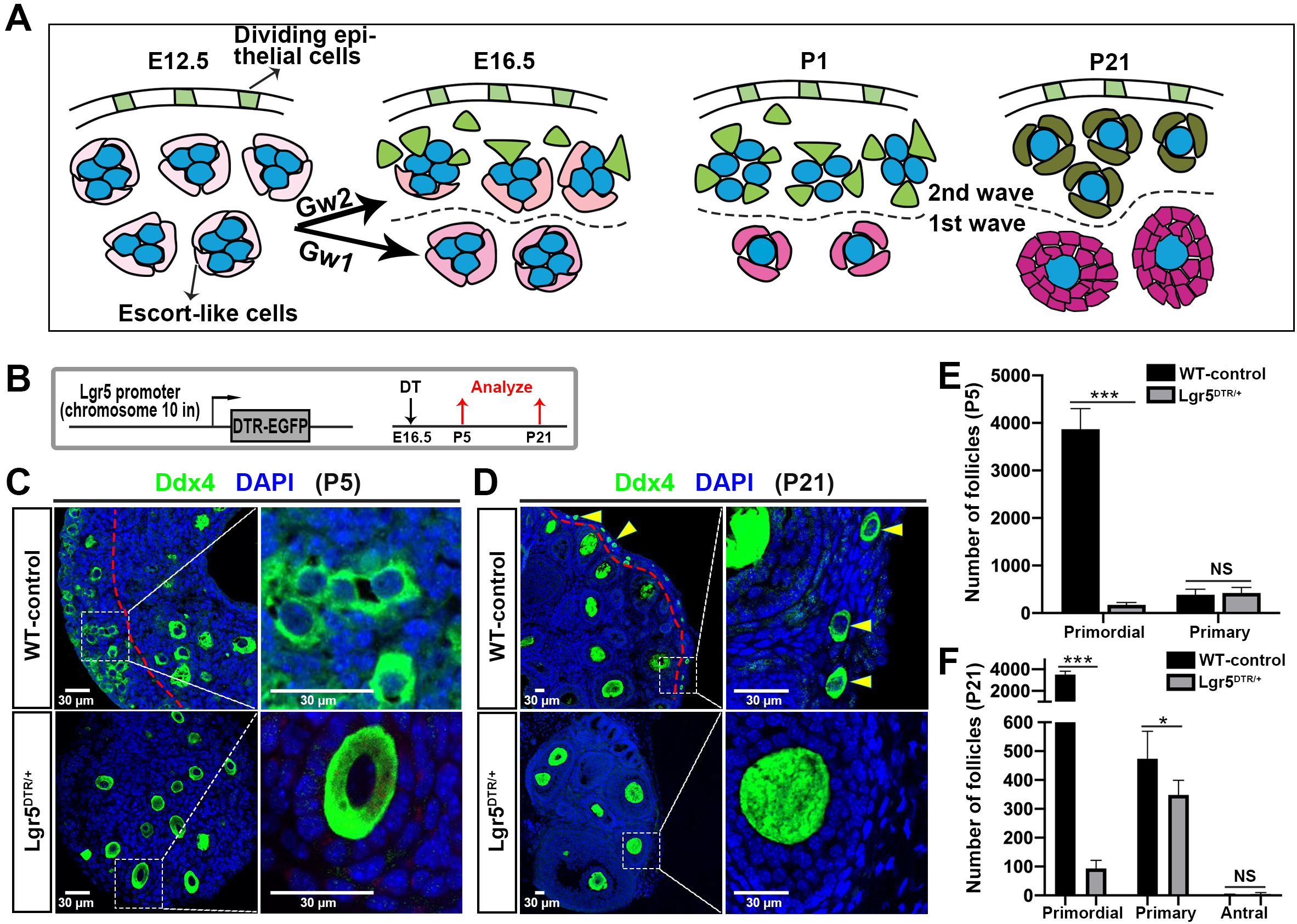
Lgr5-expressing cell ablation impairs the second wave follicle formation. (A) Model of distinct roles of escort-like cells (ELCs, pink) and surface-derived Lgr5^+^ epithelial pregranulosa cells (green) in forming the two waves of follicles. (B) Experimental strategy to ablate Lgr5-expressing cells using the Lgr5-DTR-EGFP mouse model. (C) Histological analysis of the ovary from wild-type mice and Lgr5^DTR/+^ animals at P5. (D) Histological analysis of the ovary from wildtype mice and Lgr5^DTR/+^ animals at P21. (E) Follicle quantification in the ovary at P5 after DT administration at E16.5. (F) Follicle quantification in the ovary at P21 after DT administration at E16.5. NS, Not significant. *P<0.05, ***P<0.001 (t-test). Scale bars: 30 μm.

### Depletion experiments confirm that Lgr5+ cells give rise mainly to the second wave of follicles

We used the Lgr5-DTR-EGFP mice (Tian et al. 2011) to ablate Lgr5^+^ cells during fetal follicle development by treatment with diphtheria toxin (DT) to test the prediction of our studies that only second wave follicles should be affected (Figure 6B). In controls which were treated with DT at E16.5, but lacked the transgene, a robust population of nearly 4,000 primordial follicles was observed at P5 in the cortical region, and about 400 rapidly developing first wave medullary primary follicles (Figure 6C, 6E). In contrast, pregnant females carrying the construct that were DT treated at E16.5 and examined at P5, contained less than 200 primordial follicles, but again about 400 first wave primary follicles (Figure 6C, 6E). Similarly, at P21, DT-treated controls contained more than 3,500 cortically located primordial follicles (Figure 6D, arrowheads) and 450 medullar primary follicles (Figure 6D, 6F). By comparison, Lgr5^DTR/+^ animals treated with DT retained less than 100 primordial follicles or about 3% of controls (Figure 6D, 6F). The number of wave one primary follicles was reduced by only about 33% to about 300. These results strongly support the conclusions of our previous experiments that Lgr5^+^ pregranulosa cells nourish second wave of follicles, but that most first wave follicles retain ELCs that differentiate and support follicle development without any contribution from surface-derived Lgr5+ cells.

## Discussion

### A gene expression roadmap for understanding mammalian fetal ovarian development and follicle formation

Single cell RNA sequencing provides a powerful approach for analyzing the cell types and active genes that mold tissue development. By generating single cell transcriptomes from more than 24,000 cells isolated from E12.5 to P5 mouse ovaries, we gained significant new insight into the cellular and genetic foundations of oogenesis. At any given age, ovarian germ cells vary by more than 24 hours in developmental stage, due to PGC asynchrony and meiotic initiation in an anterior to posterior wave during this period (Arora et al., 2016; Bullejos and Koopman, 2004; Menke et al., 2003). However, by reconstructing developmental trajectories based on gene expression we showed that data from germ cells aligned on a single pseudotimeline of cellular and meiotic development (Figure 1D).

Our studies of somatic cells revealed three main subgroups, containing at least eighteen somatic cell clusters, which probably include most major cell types. By re-analyzing the seven clusters within the “granulosa” subgroup we obtained fourteen clusters that represent an even higher resolution description of cell types or cell states that occur during ovarian and follicle development up to P5. The presence of multiple distinct clusters of very similar cells at different times argues that like germ cells, multiple somatic cell types follow timelines of changing gene expression. The data underlying this study will be useful for many subsequent studies of ovarian development, including cyst development and fragmentation, meiotic induction and progression, nurse cell-oocyte selection, follicle wave specification and other developmental processes.

### An initial population of escort-like pregranulosa cells mediates germline-soma interactions

Despite this potentially broad utility, we focused on one important problem requiring a detailed analysis of ovarian somatic cells, namely the origin and development of the two follicular waves. First, we identified somatic escort-like cells, a large, initially uniform population that we profiled at E12.5. ELCs interact with all incoming germ cells in both sexes based on histology (Pepling and Spradling, 2001; Fig. 3J). We propose that ELCs correspond to bipotential cellular precursors (McLaren, 1991; Mork et al. 2012a), as they initially surround both female and male germline cysts (Fig. 3E, 3F, 3J). The relatively high expression of Wnt4 and Wnt6 and of BMP2 in ELCs in E12.5 ovaries suggests that they participate in Wnt signaling-dependent early female differentiation pathways. Our observations support the idea that female germ cell development starts out uniformly, and that differences underlying the two follicular waves only arise later.

True escort cells (ECs), in Drosophila, comprise a squamous epithelial cell type with many similarities to ELCs. ECs or their earlier precursors known as “intermingled cells” (Gilboa and Lehmann, 2007; Slaidina et al., 2019), interact with PGCs prior to adulthood and with new germ cells generated by stem cell division (Morris and Spradling, 2011). In particular, ECs surround and signal to germ cells during stages of cyst formation and meiosis analogous to those that take place in the mouse ovary between E10.5 and E18.5. If germ cells at these stages are ablated in adult Drosophila ovaries, ECs turn over (Kai and Spradling, 2003). Wnt signaling from escort cells interacts antagonistically with terminal BMP signaling to establish a gradient in escort cells that is important for ongoing germ cell development (Song and Xie, 2004; Wang et al. 2015a; Mottier-Pavie et al. 2016; Wang and Page-McCaw, 2018). In addition, disrupting EC gap junctions (Mukai et al. 2011) or steroid signaling (Morris and Spradling, 2012) in escort cells arrests germ cell development. The transcriptomes of mouse ELCs (Table S3) will make it easier to uncover potential parallel mechanisms utilized by ELCs and ECs during early stages of germ cell development.

### Surface-derived Lgr5-positive pregranulosa cells displace ELCs in the ovarian cortex but not in the medulla

A second population of pregranulosa cells arises after E12.5 from dividing progenitors in the ovarian surface epithelium. Daughters invade into the cortical layers of the ovary and interact with developing cysts and their associated ELCs. The production of these surface-derived pregranulosa cells appears to fall sharply after E18.5 (Figure 4D), and despite ongoing cell division by Lgr5+ cells in the surface layer, new pregranulosa cells were not being generated by P1. Our experiments show that surface-derived pregranulosa cells quickly associate with developing germ cells. Gng13 expression, which is much higher in surface-derived cells than in ELCs, reveals that surface-derived cells have penetrated into the most superficial cortical layers by E14.5 (Figure 4E), although they do not yet fully surround germ clusters even near the surface. However, lineage labeling of ELCs shows that at E12.5 they are found associated with germ cells throughout the ovary, while by E19.5 they only remain around germ cells in the medullar region. Labeling using Lgr5-cre reveals a complementary pattern where these cells fully surround cortical germ cells by P1.

The surface granulosa cells invade the cortical follicles at a time when germ cells are undergoing germline cyst fragmentation (Lei and Spradling, 2013; Lei and Spradling, 2016). Arriving pregranulosa cells might contribute to intercellular bridge breakage, by forcing out ELCs in a way that damages some intercellular bridges, which they often encircle. However, the rates of clonal labeling, cyst formation and initial fragmentation of cortical and medullar cysts are indistinguishable (Lei and Spradling, 2013), probably reflecting the continuing influence of ELCs. After birth, medullar cysts complete breakdown faster, consistent with their accelerated development into growing follicles. Thus, the available evidence argues against the idea that invading Lgr5+ pregranulosa cells influence cyst breakdown.

### First and second wave follicles utilize different granulosa cells

The developmental trajectory analysis revealed that two subclasses of pregranulosa/granulosa cells have been generated by E18.5. Lineage analysis and gene expression verified that these correspond to Gw1 cells on first wave follicles that derive from ELCs, and Gw2 cells on second wave follicles that derive from surface progenitors. We propose (Figure 6A) that all germ cells undergo early development in contact with a relatively uniform population of ELCs. After E12.5, first wave follicles continue to develop through interactions with ELCs, which simultaneously differentiate into Gw1 pregranulosa and granulosa cells. In contrast, in the cortical regions, the influence of ELCs declines as surface-derived pregranulosa cells fully replace them by E19.5. From this time forward, wave 2 follicles develop exclusively through interactions with Gw2 granulosa cells. As a test of this model we ablated Lgr5-expressing cells at E16.5 using Lgr5-DTR-EGFP and diphtheria toxin treatment (Tian et al. 2011). As predicted, first wave follicles were scarcely affected, but 2nd wave follicles were almost completely eliminated, showing their sensitivity to loss of Lgr5-expressing somatic cells.

### What is the significance of somatic cell differences for first and second wave follicles?

Our data provide new information on the biological contributions of first wave follicles to fertility that are in agreement with the results of (Zheng et al. 2014), and contradict the view that first wave follicles are lost to atresia before contributing to fertility (McGee et al., 1998). Moreover, the Lgr5-DTR-EGFP mice treated with DT contained follicles derived from the first wave and were fertile.

Our observation that first and second wave follicles are nurtured by granulosa cell populations with different patterns of gene expression might reflect several mechanisms, including known differences in the rate of development, the onset of quiescence and the activation of follicle growth and maturation in these two follicle populations. Wave one follicles develop faster than wave two follicles, do not enter quiescence, and rapidly embark as primary follicles on an accelerated maturation program to allow fertility by the time of puberty. The rate of follicle development in several organisms can be strongly modulated by nutrition as reflected in insulin signaling (Laws and Drummond-Barbosa, 2017), and mammalian follicular growth is also influenced by activin/inhibin and steroid signaling (Myers et al. 2009; Ojima et al. 2019). Several gene changes selective for Gw1 cells by P1 are candidates for mediating a wave-specific effect on growth, including Cdn1c, Hsd17b11, Hsd17b7, Hsd17b1, Hsd3b1, Rap2b, Hmgcs2 and Thbs1 (Figure S4). In contrast, wave two follicles cease proliferation around P5 and become quiescent primordial follicles. Gw2 selective genes that might influence wave two follicle development include Lgr5, Lhx9, Klf2, Gng13, and Aldh1a2.

Further studies will be required to learn the functional roles these genes play on first or second wave follicles. When primordial follicles leave quiescence and undergo maturation, their granulosa cells may express the same genes that appear to be specific for Gw1 cells in P1 to P5 animals. It remains unclear if any of the differences between Gw1 and Gw2 gene expression reflect more than timing differences or genes involved in the induction and release from primordial follicle quiescence. However, our studies leave open the possibility that there are biological differences between the oocytes of very young mothers (derived from wave 1 follicles) compared to those from older mothers, regulating from granulosa cell expression differences.

### Primordial follicle pool size might depend on the level of surface granulosa cell production

If differences in granulosa cell gene expression program the two follicular waves, then it might be possible to alter their proportions by manipulating the number of surface-derived pregranulosa cells. Treatments that increase the number of these cells, for example, by increasing the number of Lgr5+ pre-granulosa-generating divisions, might cause more cells to invade the ovary and displace ELCs from a larger fraction of cysts and thereby increase the size of the primordial follicle pool. There would be a corresponding reduction of wave 1 follicles, with unknown consequences.

### Evolutionary conservation of somatic cell behavior during follicle formation

Many aspects of early female germline development and follicle formation are known to be highly conserved in evolution. In many organisms, germ cells initially form germline cysts that differentiate into both oocytes and nurse cells (see Büning, 1994), and the biology of meiosis and recombination are highly conserved. Our study provides evidence that somatic cells likewise play conserved roles during cyst development and follicle formation. Mouse escort-like cells have striking similarities to Drosophila escort cells in their interactions with developing cysts and gene expression. We show here that escort like cells are normally replaced by Lgr5+ pre-granulosa cells that migrate in from the ovarian cortex during fetal stages. The same replacement takes place in Drosophila for all cysts derived from germline stem cells. In this case, the replacing cells are the daughters of somatic stem cells located on the surface of each ovariole. Moreover, in Drosophila, like the mouse, the first few follicles are made from germ cells derived directly from primordial germ cells (and not from stem cells) (Asaoka and Lin, 2004). These first germ cells are wrapped by escort cells that are probably not replaced since follicle cell stem cells have not yet appeared. These initial Drosophila follicles are also notable in developing faster than later follicles derived from stem cells. Thus, the first 100 or so follicles produced by young females have follicle/granulosa cells that rapidly differentiated using only escort cell precursors, a striking parallel to the wave 1 population in the mouse. The finding that the strategy of producing two groups of follicles with distinctive somatic supports appear to have been preserved throughout animal evolution reveals again the value of an evolutionary comparative approach to understanding tissue development and function.

## ACKNOWLEDGMENTS

We are grateful to the Johns Hopkins University School of Medicine Biotechnology center for assistance with some of the scRNAseq experiments. We especially thank Allison Pindar and Fred Tan of the Carnegie Embryology Biotechnology Center for assistance in carrying out scRNAseq and in data analysis. We are grateful to Mike Sepanski for carrying out electron microscopy. We thank Dr. Frederic J. de Sauvage (Genentech, Inc.) for kindly providing us with Lgr5-DTR-EGFP mice.

## AUTHOR CONTRIBUTIONS

W.N. and A.C.S. designed experiments, analyzed data and wrote the manuscript. W.N. performed research.

## Experimental Methods

### Animals

Mouse experiments in this study were performed in accordance with protocols approved by the Institutional Animal Care and Use Committee (IACUC) of the Carnegie Institution of Washington. Lgr5-DTR-EGFP mice were obtained from Genentech (South San Francisco, CA). R26R-tdTomato mice were obtained from Chen-Ming Fan lab (Carnegie Institution for Science, MD). Lgr5-CreERT2 mice (008875), Axin2-CreERT2 mice (018867) and R26R-EYFP reporter mice (006148) were acquired from the Jackson Laboratory.

### Labeling and Tracing Experiments

The R26R-tdTomato females were crossed with the Lgr5-CreERT2 males, those with a vaginal plug were considered as E0.5. The pregnant females at E13.5 or newborn pups at P1 were given a single intraperitoneal (i.p.) injection of tamoxifen (Tmx; 10 mg/ml in corn oil) at 1 mg per 25 g body weight. The R26R-EYFP females were crossed with the Axin2-CreERT2 males, and the pregnant females at E10.5 were injected i.p. with tamoxifen at 0.2 mg per 25 g body weight.

### Diphtheria Toxin injection

Pregnant mice (E16.5) were injected i.p. with 10 μg/kg of diphtheria-toxin solution in PBS.

### Immunofluorescence and immunohistochemistry

Ovaries were fixed in cold 4% Paraformaldehyde overnight, incubated sequentially in 10% and 20% sucrose in PBS overnight, embedded in OCT, and stored at −80°C until cryosectioning. After high-temperature antigen retrieval with 0.01% sodium citrate buffer (pH 6.0), the frozen sections (10 μm) were blocked with 10% normal donkey serum for 30 mins, and then incubated with primary antibodies overnight at 4°C. The primary antibodies used are presented in KEY RESOURCES TABLE. For immunofluorescence, the sections were washed with wash buffer and incubated with the appropriate Alexa-Fluor-conjugated secondary antibodies (1:200, Invitrogen) at room temperature for 2 hours. After staining with DAPI, samples were analyzed using confocal microscopy (Leica SP5). For immunohistochemistry, the slides were incubated with avidin-conjugated secondary antibodies (ab64264, Abcam) before being exposed to diaminobenzidine (DAB, ab64264, Abcam) for 1 min and then counterstained with hematoxylin.

### In situ hybridization

Tissue samples were fixed in neutral buffered formalin (NBF 10%) at room temperature for 24 hrs, embedded in paraffin, and sectioned to a thickness of 5 μm. Tissue was pretreated with boiling 1X Target Retrieval followed by Protease III at 40°C for 30 mins. After pretreatment, the samples were hybridized with probes against mouse *Wnt6* (401111), *Lgr5* (312171-C2), *Gng13* (462531), and *Fmr1* (496391-C2) using RNAscope 2.5 HD Duplex Assay (322430, ACDBio). Signal was detected by two different chromogenic substrates (HRP-C1-Green and AP-C2-Red). Finally, slides were counterstained with hematoxylin and covered with mounting medium.

### Tissue dissociation and single cell library preparation

Perinatal ovaries were dissected and placed in 1X PBS on ice, then dissociated into single cells using 0.25% Trypsin at 37°C with pipet trituration at intervals. E12.5 ovaries were dissociated for 20 min, E14.5 ovaries for 40 min, E18.5 ovaries for 1h, P1 and P5 ovaries for 80 min. After neutralized with 10% FBS, dissociated cells were passed through 70 μm and 30 μm cell strainer, separately. Approximately 10,000 live cells were loaded per sample onto the 10x Genomics Chromium Single Cell system using the v2 chemistry per manufacturer’s instructions (Zheng, et al., 2017). Single cell RNA capture and library preparations were performed according to manufacturer’s instructions. Sample libraries were sequenced on the NextSeq 500 (Illumina). Sequencing output was processed through the Cell Ranger 2.1.0 mkfastq and count pipelines using default parameters. Reads were quantified using the mouse reference index provided by 10x Genomics (refdata-cellranger-mm10v1.2.0). 10X genomics chemistry version 3’c2; mouse transcriptome mm10, cell ranger 2.1.1.

### Cell identification and clustering analysis

Package “Seurat” v2.3.4 (https://satijalab.org) (Butler, et al., 2018) was used to analyze the single cell RNA-seq data. The count data produced by Cell Ranger pipelines was read and transformed into Seurat object using the *CreateSeuratObject* function. Cells with too few reads were filtered out using *FilterCells* function (subset.names = “nGene”, low.thresholds = 500). Filtered count matrices for each library (E12.5, E14.5, E18.5, P1 and P5) were Log-normalized, scaled, and merged to integrated dataset through *MergeSeurat* function. After detecting the variable genes (x.low.cutoff = 0.0125, x.high.cutoff = 5, y.cutoff = 0.5), cell clusters were determined and identified based on the SNN algorithm (reduction.type = “pca”, dims.use = 1:10, resolution = 0.6), and visualized through dimensionality reduction by *RunTSNE* function. For reanalyzing germ population or granulosa population, clusters with unique expression of germ cell markers or granulosa cell markers were extracted from integrated dataset by *SubsetData* function. The isolated cluster was divided into several subclusters after a series of normalization, scale and dimensionality reduction.

### Cell trajectory analysis

The epithelial cells, escort-like cells and pregranulosa cells at E12.5, E14.5 and E18.5 were isolated from the integrated granulosa dataset (Figure 3C) using *SubsetData* function in Seurat package. Then we used the monocle2 (Qiu, et al., 2017) to reconstruct the differentiation pathways across the above cell types. The Seurat object was converted to CellDataSet object through *importCDS* function. We used *differentialGeneTest* (fullModelFormulaStr = “∼Cluster”) to test differential expression as a function of pseudotime, and used the *reduceDimension* function to run the DDRTree algorithm and estimate the ordering of cells along a trajectory. Subsequently, we selected cells that were placed on the Gw1 or Gw2 branch (Figure 4G), and compared the differential expression between Gw1 and Gw2 at each developmental stage through *VlnPlot* function (Figure S4). For P1 and P5 dataset, the same strategy was performed to run the pseudotime analysis.

## QUANTIFICATION AND STATISTICAL ANALYSIS

The significance of experimental treatments was analyzed by Students t-test (single-tailed). Significance levels are shown on the Figures using the following code: ∗ = p<0.05; ∗∗ = p<0.01; ∗∗∗ = p<0.001.

## MATERIAS, DATA AND SOFTWARE AVAILABILITY

The raw and processed data files are available in GEO: GSE136441

Further information and requests for resources and reagents should be directed to and will be fulfilled by the corresponding author, Allan Spradling.

