## Supplemental Figures and Table. for "Escort-like somatic cells mediate early mouse fetal ovarian development but surface-derived Lgr5+ cells support primordial follicles"

### Supplemental Material

#### Supplemental Tables

Table S1. Single cell RNA seq parameters

| Stage | Number of Cells | UMI/cell | genes/cell |
|---|---|---|---|
| E12.5 | 4,081 | 12,093 | 3,399 |
| E14.5 | 9,004 | 7,258 | 2,483 |
| E18.5 | 3,073 | 8,351 | 2,843 |
| P1 | 4,708 | 8,291 | 2,583 |
| P5 | 3,362 | 8,864 | 2,281 |

UMI/cell and genes/cell represent averages across the all cell types, and may not be comparable to corresponding values for restricted or single cell groups in Tables S2-S4.

Table S2 Transcriptomes for cell clusters in Figure 2C

Gene, total UMIs/cell, mean UMIs/cell are shown for each cluster: Premeiotic, Pre-leptotene (Pre-L), Leptotene, Zygotene, Pachytene, Diplotene, Late Diplotene, Dictyate.

Table S3 Transcriptomes for cell clusters in Figure 3C

Gene, total UMIs/cell, mean UMIs/cell are shown for each cluster: Escort_1, Escort_2, Escort_3, Epithelial_1, Epithelial_2, Epithelial_3, Epithelial_4, Pregranulosa_1, Pregranulosa_2, Pregranulosa_3, Granulosa_1, Granulosa_2, Granulosa_3, Granulosa_4, Outliers_1, Outliers_2. Note that Escort_1, Escort_2 and Escort_3 correspond to ELCs.

Table S4 Transcriptomes for Gw1 and Gw2

Gene, Gw1 UMI/cell, Gw2 UMI/cell, Gw1/Gw2 fold change (log2) are shown at each time point: E12.5, E14.5, E18.5, P1 and P5.

**Supplemental Figures**

**Figure S1. Expression pattern of mesenchymal cells in perinatal ovaries.**

(A) Cellular localization of Nr2f2 in perinatal ovaries. Ovaries were stained for N2rf2, Foxl2, and the oocyte marker DDX4 at E12.5, and E18.5. (B) Cellular localization of Col1a1 in perinatal ovaries. Ovaries were stained for Col1a1, and the oocyte marker DDX4 at E14.5, and P2. DAPI was used to identify nuclear DNA.

**Figure S2. Heat map reports differential expression of meiosis related gene sets for each meiosis cluster.**

(A) Heat map depicting differentially expressed genes in each meiosis cluster compared to all other clusters. Colored bars represent UMI expression levels of individual cells to provide information on observed variation in expression, including statistical noise. Specific meiosis related genes used to annotate clusters are indicated to the right of the heat map. Color scheme is based on z-score distribution from -2 (blue) to 2 (orange).

**Figure S3. Expression pattern of Wnt and Bmp pathway components in escort-like cells.**

(A) The relative expression in Escort-like cells of different Wnt ligands in E12.5 (salmon) and E14.5 (green) ovaries. (B) The relative expression in Escort-like cells of different Bmp pathway components in E12.5 (salmon) and E14.5 (green) ovaries. (A) The violin plots show the top differentially expressed genes in Gw1 cells compared to Gw2. Developmental state was separated by dashed line. y-axis scale: a normalized UMI/cell scale for each gene to facilitate display.

**Figure S4. Gene profiles comparison shows the different development between Gw1 and Gw2 pregranulosa cells.**

(A) The violin plots show the top differentially expressed genes in Gw1 cells compared to Gw2. Developmental state was separated by dashed line. (B) The violin plots show the top differentially expressed genes in Gw2 cells compared to Gw1. Developmental state was separated by dashed line. (C) The expression pattern of markers that chart the progression of Gw1 cells (from E14.5 to P5). (D) The expression pattern of markers that chart the progression of Gw2 cells (from E14.5 to P5). y-axis scale: a normalized UMI/cell scale for each gene to facilitate display.

**Figure S5. Lineage tracing experiments demonstrate that epithelium derived Lgr5+ pregranulosa cells form the second wave follicles before birth but not after birth.**

(A) Schematic of the lineage tracing strategy. $Lgr5^{CreERT2/+}$ mice are crossed to $R26R^{tdT/tdT}$ reporter mice. Tamoxifen (TM) was administrated at P1, and samples were collected at P2 and P6. (B, C) Lineage tracing of Lgr5+ pregranulosa cell progeny showed that majority of tdTomato+ cells widely located in the ovary surface, whereas only a small proportion of second wave follicles with tdTomato staining.

### Fig. S1

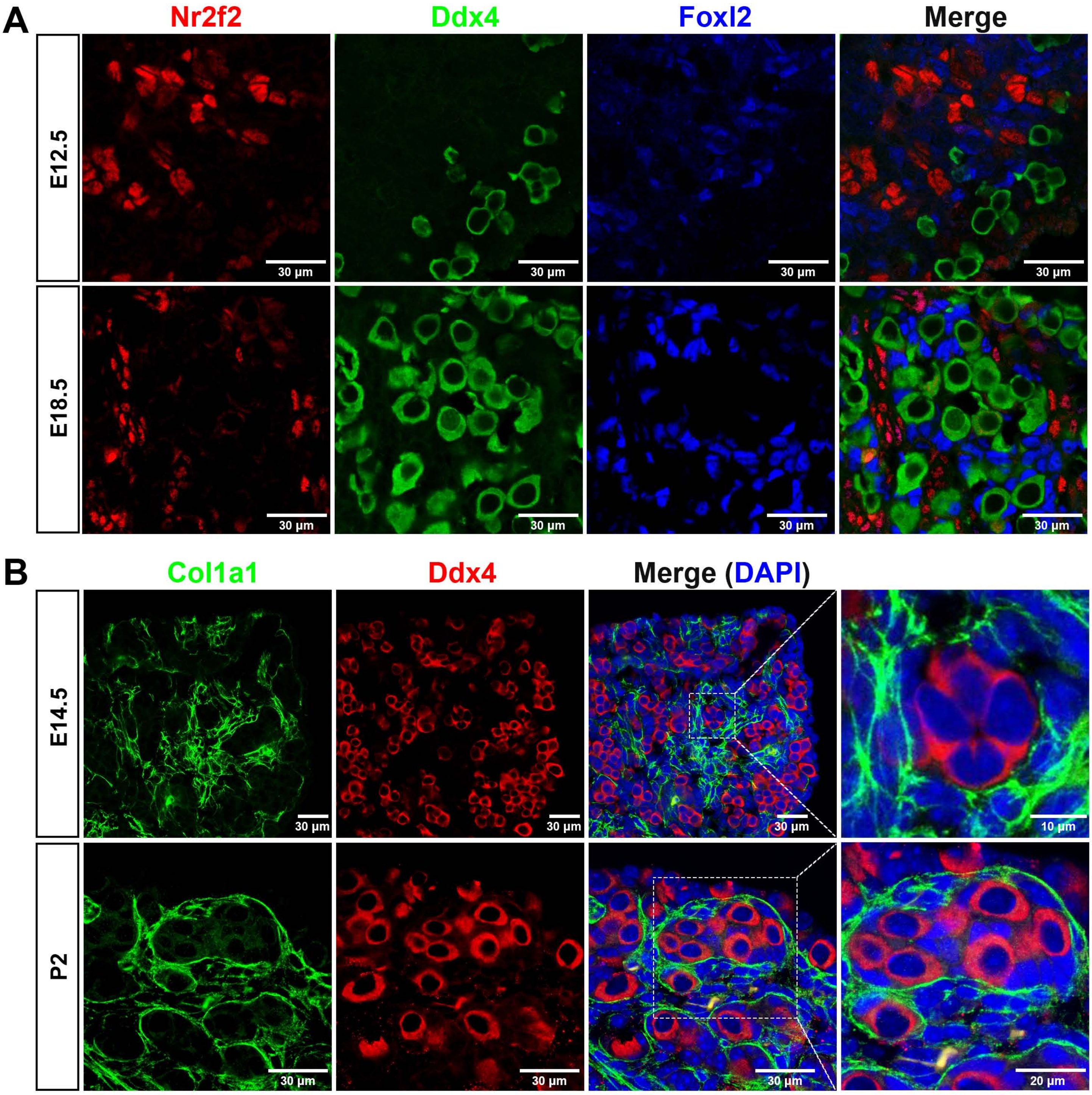
A
Nr2f2
Ddx4
Foxl2
Merge
E12.5
E18.5
30 µm
B
Col1a1
Ddx4
Merge (DAPI)
E14.5
P2
30 µm
10 µm
20 µm

Fig. S2

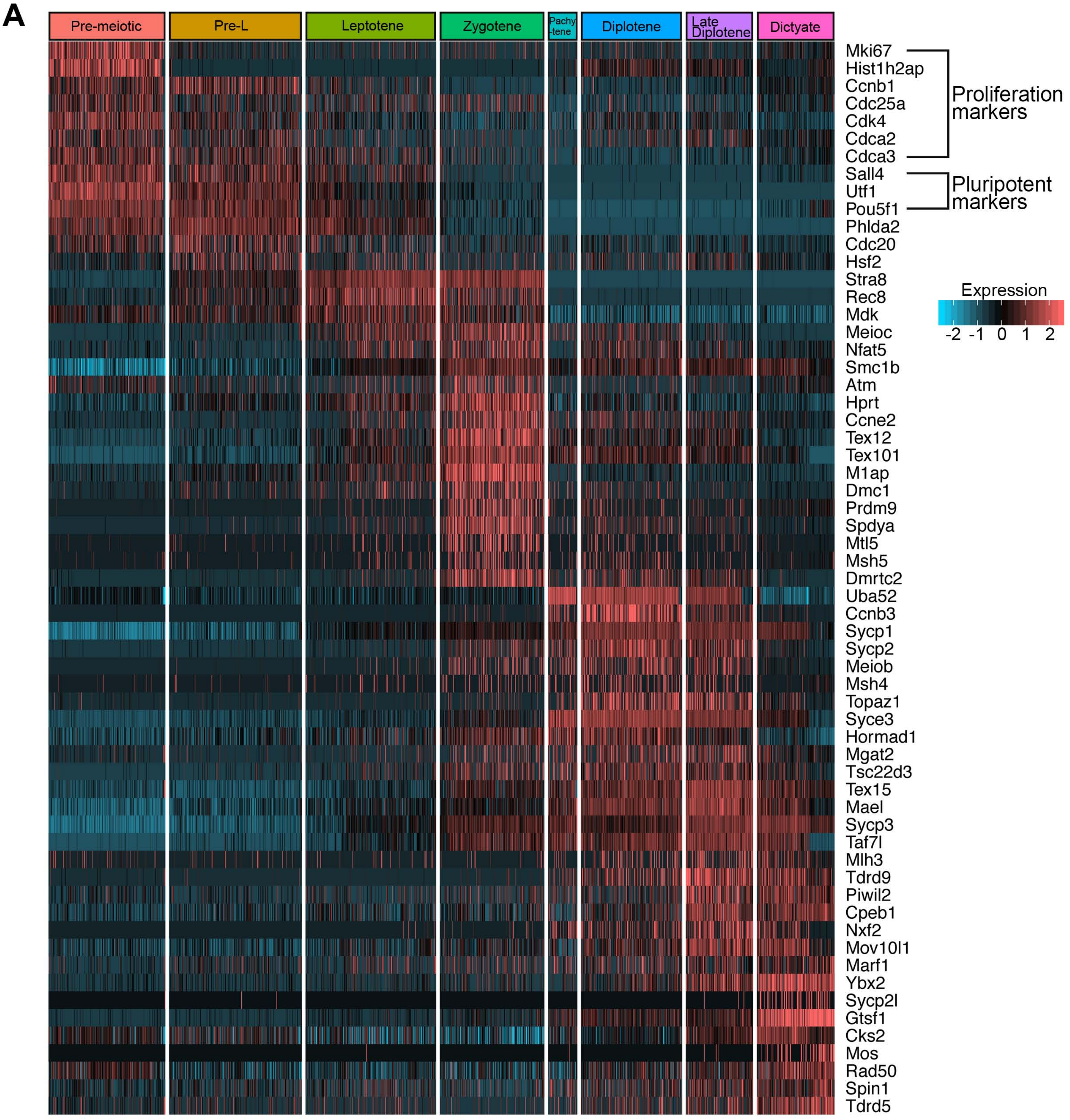
A
Pre-meiotic
Pre-L
Leptotene
Zygotene
Pachy-tene
Diplotene
Late Diplotene
Dictyate
Mki67
Hist1h2ap
Ccnb1
Cdc25a
Cdk4
Cdca2
Cdca3
Sall4
Utf1
Pou5f1
Phlda2
Cdc20
Hsf2
Stra8
Rec8
Mdk
Meioc
Nfat5
Smc1b
Atm
Hprt
Ccne2
Tex12
Tex101
M1ap
Dmc1
Prdm9
Spdya
Mtl5
Msh5
Dmrtc2
Uba52
Ccnb3
Sycp1
Sycp2
Meiob
Msh4
Topaz1
Syce3
Hormad1
Mgat2
Tsc22d3
Tex15
Mael
Sycp3
Taf7l
Mlh3
Tdrd9
Piwil2
Cpeb1
Nxf2
Mov10l1
Marf1
Ybx2
Sycp2l
Gtsf1
Cks2
Mos
Rad50
Spin1
Tdrd5
Proliferation markers
Pluripotent markers
Expression
-2 -1 0 1 2

### Fig. S3

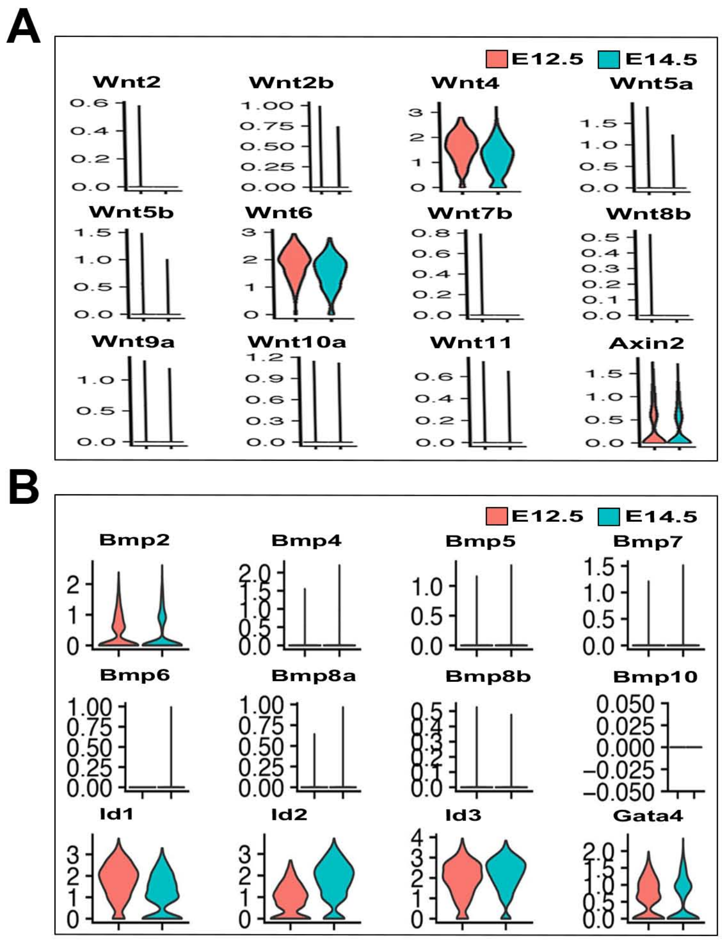
A
E12.5
E14.5
Wnt2
Wnt2b
Wnt4
Wnt5a
Wnt5b
Wnt6
Wnt7b
Wnt8b
Wnt9a
Wnt10a
Wnt11
Axin2
B
E12.5
E14.5
Bmp2
Bmp4
Bmp5
Bmp7
Bmp6
Bmp8a
Bmp8b
Bmp10
Id1
Id2
Id3
Gata4

Fig. S4

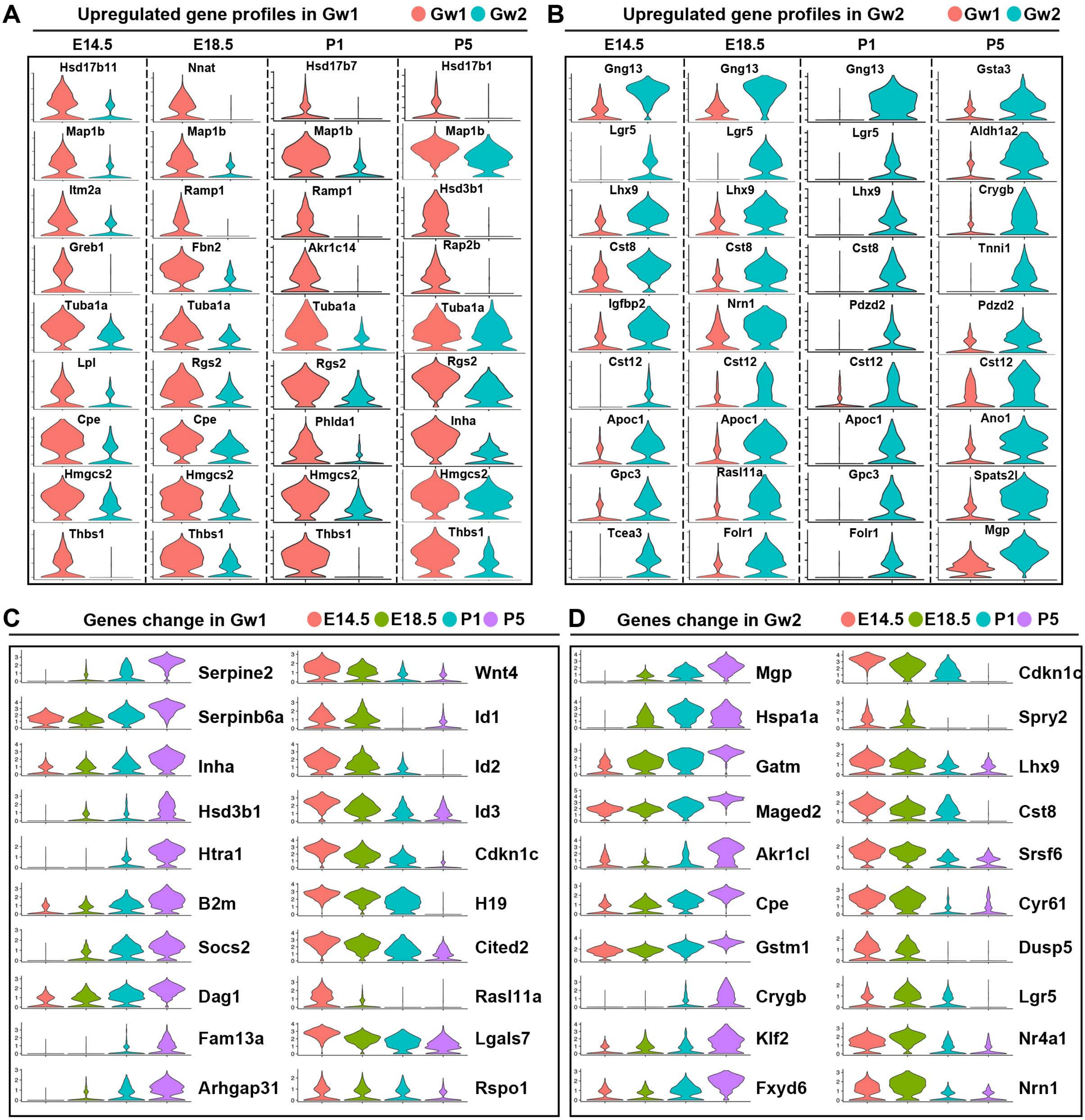
A
Upregulated gene profiles in Gw1
Gw1 Gw2
E14.5
E18.5
P1
P5
Hsd17b11
Nnat
Hsd17b7
Hsd17b1
Map1b
Itm2a
Ramp1
Hsd3b1
Greb1
Fbn2
Akr1c14
Rap2b
Tuba1a
Lpl
Rgs2
Cpe
Phlda1
Inha
Hmgcs2
Thbs1
B
Upregulated gene profiles in Gw2
Gng13
Gsta3
Lgr5
Aldh1a2
Lhx9
Crygb
Cst8
Tnni1
Igfbp2
Nrn1
Pdzd2
Cst12
Apoc1
Ano1
Gpc3
Rasl11a
Spats2l
Tcea3
Folr1
Mgp
C
Genes change in Gw1
E14.5 E18.5 P1 P5
Serpine2
Serpinb6a
Inha
Hsd3b1
Htra1
B2m
Socs2
Dag1
Fam13a
Arhgap31
Wnt4
Id1
Id2
Id3
Cdkn1c
H19
Cited2
Rasl11a
Lgals7
Rspo1
D
Genes change in Gw2
Mgp
Hspa1a
Gatm
Maged2
Akr1cl
Cpe
Gstm1
Crygb
Klf2
Fxyd6
Cdkn1c
Spry2
Lhx9
Cst8
Srsf6
Cyr61
Dusp5
Lgr5
Nr4a1
Nrn1

### Fig. S5

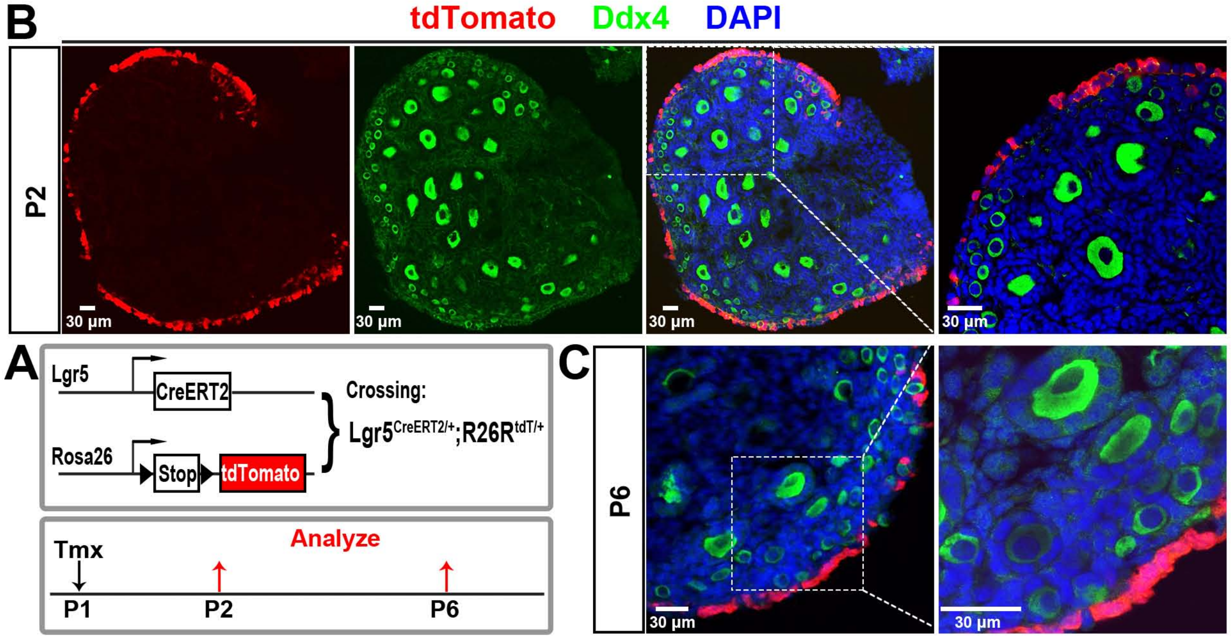
B
tdTomato Ddx4 DAPI
P2
30 µm
30 µm
30 µm
30 µm
A
Lgr5
CreERT2
Rosa26
Stop
tdTomato
Crossing:
Lgr5CreERT2/+;R26RtdT/+
Tmx
Analyze
P1
P2
P6
C
P6
30 µm
30 µm
